# GARFIELD-NGS: Genomic vARiants FIltering by dEep Learning moDels in NGS

**DOI:** 10.1101/149146

**Authors:** Viola Ravasio, Marco Ritelli, Andrea Legati, Edoardo Giacopuzzi

## Abstract

**Summary:** Exome sequencing approach is extensively used in research and diagnostic laboratories to discover pathological variants and study genetic architecture of human diseases. However, a significant proportion of identified genetic variants are actually false positive calls, and this pose serious challenges for variants interpretation. Here, we propose a new tool named GARFIELD-NGS (Genomic vARiants FIltering by dEep Learning moDels in NGS), which rely on deep learning models to dissect false and true variants in exome sequencing experiments performed with Illumina or ION platforms. GARFIELD-NGS showed strong performances for both SNP and INDEL variants (AUC 0.71 - 0.98) and outperformed established hard filters. The method is robust also at low coverage down to 30X and can be applied on data generated with the recent Illumina two-colour chemistry. GARFIELD-NGS processes standard VCF file and produces a regular VCF output. Thus, it can be easily integrated in existing analysis pipeline, allowing application of different thresholds based on desired level of sensitivity and specificity.

**Availability:** GARFIELD-NGS available at https://github.com/gedoardo83/GARFIELD-NGS

## Introduction

DNA analysis through exome sequencing is now the main tool to discover disease related variants (Koboldt *et al.*, 2013; Wang *et al.*, 2013). However, variants identified by exome sequencing often carries a significant proportion of false positive calls, especially INDELs, and this pose serious challenges for variants interpretation (Zhang *et al.*, 2015; Jiang *et al.*, 2015; Damiati *et al.*, 2016). Advanced methods based on machine learning have been developed for large datasets, while few effective solutions are available for small experiments. Here, we propose a new tool, Genomic vARiants FIltering by dEep Learning moDels in NGS (GARFIELD-NGS), that rely on deep learning models to effectively classify true and false variants in exome sequencing experiments performed on both Illumina or ION platforms.

## Methods

Starting from 23 high-coverage exome sequencing experiments on NA12878 reference sample, we assembled two pools of 178,450 Illumina variants (173,116 SNVs / 5,334 INS/DELs) and 181,479 ION variants (177,362 SNVs / 4,117 INS/DELs). True and false variants were determined based on the comparison with NA12878 high confidence calls from NIST v.3.3.2 (Zook *et al.*, 2014). Variants in each group were splitted randomly in 4 independent datasets (pre-training, training, validation and test). Additional 60X and 30X test sets were produced by random subsampling of the original sequencing data, while HiSeqX test set was based on 3 experiments produced on HiSeq X platform. We evaluated 18 features for ION variants and 10 for Illumina variants (Supplementary Table S1) to generate 4 distinct prediction models based on multilayer perceptron algorithm as implemented in H2O v.3.10.4.5 deep learning method (http://www.h2o.ai): Illumina INS/DELs, Illumina SNVs, ION INS/DELs, and ION SNVs. After hyper-parameters optimization using random search, performances of the final models were assessed on test sets and validated on the replication sets, composed by 4 additional experiments not used in model development. GARFIELD-NGS performances on test and replication sets were compared to well established hard-filters, including GATK VQSR method for Illumina data (Van der Auwera *et al.*, 2013) and previously published hard-filters for ION data (Damiati *et al.*, 2016). Finally, we assessed how our models filter variants from data not processed by our pipeline, including 35 Illumina and 32 ION WES experiments, as well as a set of 211 variants previously validated by Sanger sequencing. Detailed methods are reported in supplementary materials.

## Results

### Prediction models performances

Using H2O deep learning algorithm, we developed 4 prediction models optimized for INS/DELs and SNVs for Illumina and ION platforms (Supplementary Table S2). Our tool calculates for each variant a confidence probability (CP) ranging from 0.0 to 1.0, with higher values associated with true variants. AUROC values > 0.90 are obtained for Illumina INS/DELs, ION INS/DELs and ION SNVs, while Illumina SNVs model shows slightly reduced performances with AUROC 0.7998 (Figure 1). Accuracy is > 0.90 for all variants categories (Supplementary Table S3). GARFIELD-NGS correctly classifies more than 95% of true variants and significantly reduces false positive variants (Supplementary Fig. S1). These performances were confirmed when applying GARFIELD-NGS on the low-coverage sets (60X / 30X) and HiSeqX set (Figure 1), as well as on the replication sets (Supplementary Table S4). GARFIELD-NGS models perform well also on WES experiments not processed with our pipeline (Supplementary Fig. S2) and on a set of Sanger validated variants from real-world diagnostic setting. Here, we obtain 0.958 and 0.878 accuracy on Illumina INS/DELs and SNVs, respectively; and 0.804 and 0.955 accuracy for ION INS/DELs and SNVs, respectively (Supplementary Table S5). Additional results including models details, analysis of features contribution, detailed description of performances and characterization of filtered variants are provided in supplementary materials.

**Fig. 1.**
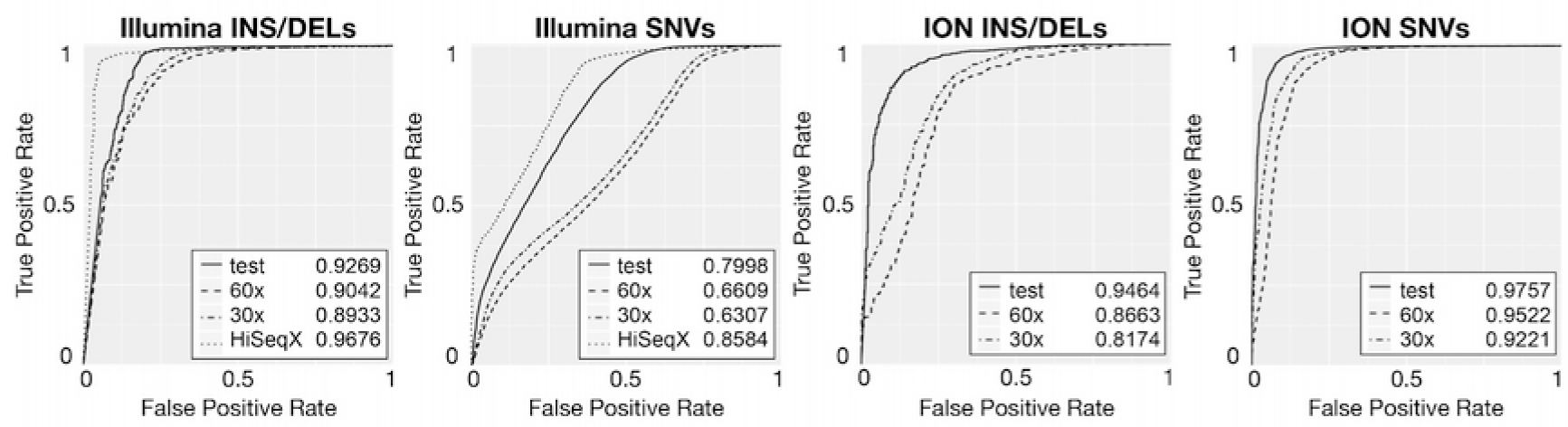
ROC curves of GARFIELD-NGS final models on test datasets. Performance of prediction models were assessed using ROC curves on test sets, 60X and 30X downsampled sets, and HiSeqX sets. Values of area under the receiver operating characteristic curve (AUROC) are indicated in the graphical plots.

### Comparison with hard-filters and VQSR

GARFIELD-NGS outperforms hard-filters in Illumina INS/DELs, ION INS/DELs and ION SNVs groups, showing higher accuracy, while it obtains comparable performances on Illumina SNVs (Supplementary Fig. S3 and Supplementary Table S3). Largest improvements are seen for INS/DELs. Accuracy of GARFIELD-NGS reaches 0.93 and 0.91 for Illumina and ION INS/DELs, respectively, compared to 0.86 and 0.80 calculated using hard-filters. When applied on INS/DELs variants GARFIELD-NGS outperforms GATK VQSR, as well. VQLOD reaches an AUROC value of 0.6783, while GARFIELD-NGS reaches 0.92 AUROC (Supplementary Fig. S4). Detailed results of performance comparisons are reported in supplementary materials.

## Discussion

Even if alternative pipelines have been proposed such as GotCloud (Jun *et al.*, 2015), SNPSVM (O’Fallon *et al.*, 2013) and DeepVariant (Poplin *et al.*, 2018), which combine variant calling and machine learning based variant filtering, the most applied variant callers for Illumina and Ion data are still GATK (Van der Auwera *et al.*, 2013; DePristo *et al.*, 2011) and TVC. Only few tools are available to directly refine SNVs and INS/DELs called using these widely adopted variant callers. GARFIELD-NGS can be applied directly to variant callers output and outperforms previous filtering strategies, obtaining robust performances even on low coverage data. The maximum accuracy thresholds retain > 95 % of true calls, while reducing false calls by 36-80 %, depending on variant category. Even at 0.99 TPR, GARFIELD-NGS maintains > 0.86 accuracy. When applied to a canonical pipeline for prioritization of disease related variants, GARFIELD-NGS significantly reduces the proportion of false candidates, thus improving identification of diagnostic relevant variants. These results define GARFIELD-NGS as a robust tool for all type of Illumina and ION exome data. GARFIELD-NGS script performs automated variant scoring on VCF files and it can be easily integrated in existing analysis pipelines.

## Funding

EG has been supported by “Fondazione Cariplo” and “Regione Lombardia” under the project Grant Emblematici Maggiori 2015-1080.

## Conflict of Interest

none declared.

